# Cell Painting predicts impact of lung cancer variants

**DOI:** 10.1101/2021.11.18.469171

**Authors:** Juan C. Caicedo, John Arevalo, Federica Piccioni, Mark-Anthony Bray, Cathy L. Hartland, Xiaoyun Wu, Angela N. Brooks, Alice H. Berger, Jesse S. Boehm, Anne E. Carpenter, Shantanu Singh

**Affiliations:** Broad Institute of Harvard and MIT; Broad Institute of MIT and Harvard; University of California, Santa Cruz; Fred Hutchinson Cancer Research Center

## Abstract

Most variants in most genes across most organisms have an unknown impact on the function of the corresponding gene. This gap in knowledge is especially acute in cancer, where clinical sequencing of tumors now routinely reveals patient-specific variants whose functional impact on the corresponding gene is unknown, impeding clinical utility. Transcriptional profiling was able to systematically distinguish these variants of unknown significance (VUS) as impactful vs. neutral in an approach called expression-based variant-impact phenotyping (eVIP). We profiled a set of lung adenocarcinoma-associated somatic variants using Cell Painting, a morphological profiling assay that captures features of cells based on microscopy using six stains of cell and organelle components. Using deep-learning-extracted features from each cell’s image, we found that cell morphological profiling (cmVIP) can predict variants’ functional impact and, particularly at the single-cell level, reveals biological insights into variants which can be explored in our public online portal. Given its low cost, convenient implementation, and single-cell resolution, cmVIP profiling therefore seems promising as an avenue for using non-gene-specific assays to systematically assess the impact of variants, including disease-associated alleles, on gene function.

## Introduction

Lung cancer is the leading cause of cancer-related mortality and presents high mutation rates ^4,5^. New variants are found every year in clinical studies, most of them Variants of Unknown Significance (VUS). Although custom-tailored assays might be created to assess the function of each gene in the presence or absence of each variant, this is exceptionally time-consuming. It is only practical for a small number of known oncogenes and tumor suppressors and is impossible for genes whose function is unknown. This limits an expansion of precision medicine, where cancer patients are tested to identify their specific mutations and ultimately receive targeted treatments.

High-dimensional profiling assays have been proposed as an accelerant for determining the significance of VUS: by measuring many phenotypic properties of cells exposed to each variant in each gene of interest, the strategy is to capture many genes’ functions in a single assay and therefore assess many variants’ impact. This strategy was successfully demonstrated using high-throughput transcriptional profiling in an approach called expression-based variant impact phenotyping (eVIP)^6,7^, where the transcriptional profiles of overexpressed reference genes (wild-type) are systematically compared to their variants (mutants) to assess impact. In this case, a bead-based, high-throughput transcriptional profiling method called L1000 was used ^8,9^.

We hypothesized that another profiling readout, image-based profiling, could also be used for variant impact phenotyping. Image-based profiling has proven powerful in more than a dozen applications in biological research and drug discovery ^10^. We sought to develop cell morphology-based Variant Impact Phenotyping (cmVIP) as a way to inexpensively assess the functional impact of coding variants for many genes using the same, systematic assay. If scaled up, a catalog might be created of all possible variants in a given oncogene or tumor suppressor to help guide clinicians.

Here, we present a systematic study of the ability of image-based profiling to characterize lung cancer variants. We conducted a high-throughput Cell Painting ^11^ experiment using gene overexpression in A549 cells to investigate the extent to which cell morphology can reveal sufficient phenotypic differences between reference genes and alleles. We developed deep learning-based computational methods to transform images of cells into high-dimensional phenotypic profiles and used them to quantify the impact of variants. In addition, we compare the performance of image-based profiling with respect to gene expression profiling to capture phenotypic changes induced by variants and to predict their functional impact.

## Results

### 1. Cell Painting captures a diversity of gene and allele phenotypes

We tested 375 overexpression perturbations (50 reference genes and 325 variants) in A549 cells using the Cell Painting assay in 384 well plates with 8 replicates each (Methods). The overexpression construct set was previously created to test the expression-based variant impact phenotyping (eVIP) method ^6^ and contains variants previously identified by exome sequencing primary lung adenocarcinomas ^1^ as well as their reference genes. They include many known impactful variants as well as many variants of unknown significance (VUS). As negative controls, we used wells with untreated cells that we call EMPTY controls.

We found that the Cell Painting assay can detect phenotypic signals for the majority of alleles (83.2%); this is an important first step in determining the impact of variants. We evaluated this as follows: after acquiring Cell Painting images for each sample (Figure 1B), we transformed them into replicate-level allele profiles using a deep learning-based workflow ^12,13^ (Figure 1A, see also Methods). We evaluated the quality of profiles using the percent replicating score ^14^, measured as the percentage of perturbations whose replicates consistently have higher similarity (reproducible signal) than random sets of perturbations; in this case 83.2% (Figure 1C).

**Figure 1.**
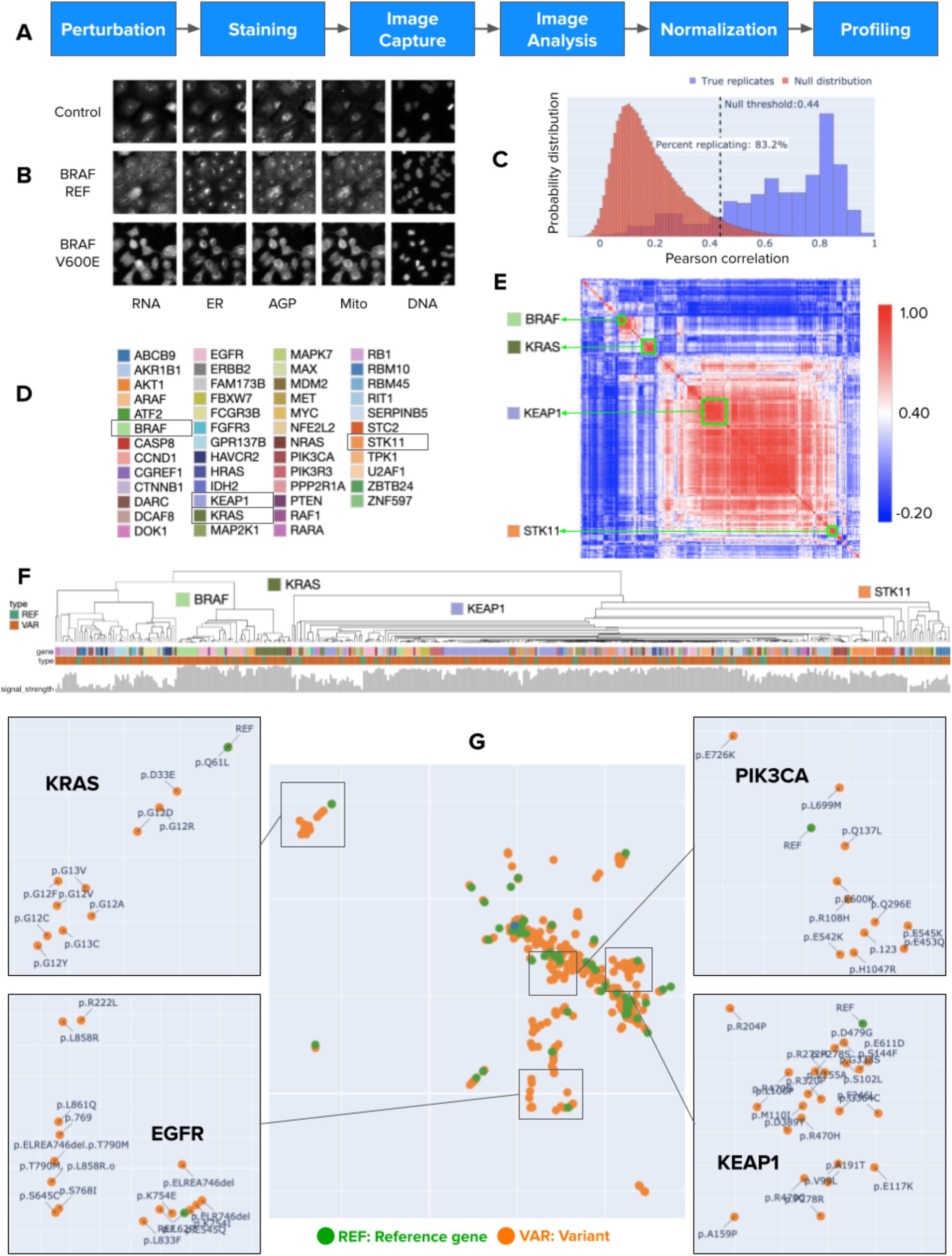
Cell morphology captures phenotypic variation of lung cancer alleles. A) Workflow to create image-based profiles by transforming Cell Painting images into quantitative, multivariate representations of the state of cells impacted by each perturbation (whether a reference gene or variant overexpressed in the cells). B) Example Cell Painting images in three experimental conditions: empty controls, BRAF reference gene overexpression, and BRAF V600E allele overexpression. The images are random crops of 200×200 pixels from a field of view (1080×1080), and each channel has been independently rescaled to fit the visible intensity range. C) Distribution of true replicates vs a null distribution of randomized replicates in this experiment, resulting in 83.2% of all perturbations having high self-correlation. Note that the null threshold (above which significant correlations are detected) is 0.44 in the Pearson correlation scale of [-1,1]. D) List of genes included in our study; some genes whose variants are grouped in the dendrogram are outlined. For each gene, we tested several variants. E) Correlation matrix between all pairs of perturbations (reference and variant overexpression) sorted according to the hierarchical clustering of the rows and columns. F) Dendrogram depicting groups found by the hierarchical clustering in the correlation matrix. The *type* bar coloring refers to whether the perturbation is a *reference* sequence or *variant*. The *gene* bar is colored according to the color code in D. G) UMAP plots of reference genes’ and variants’ perturbation-level profiles (combining data from all replicate wells). Clusters of reference genes and their variants are observed and four examples are zoomed in (Full-scale figures available at http://broad.io/cmvip/umap.html).

### 2. Variant phenotypes cluster consistently with the corresponding reference gene’s phenotype

Having determined that most reference genes and their variants’ overexpression produced a replicable profile, we next sought to assess the structure of the relationships (similarities) among those profiles. Cell Painting has been shown to recapitulate genetic pathway relationships between reference genes in overexpression perturbation experiments ^15^; here, we tested a high number of alleles per gene. After aggregating replicate-level profiles into perturbation-level profiles to obtain a high-dimensional representation of each gene and allele in our experiment, we clustered them.

The correlation matrix (Figure 1E) displays a large set of genes and alleles that have highly similar phenotypic characteristics, which indicates that within this dataset most cancer variants share the same major phenotype. Cell Painting profiles are still able to capture subtle and meaningful variations between alleles as reflected in the continuous groups of reference genes and their corresponding variants in the hierarchical clustering (Figure 1F, color bar marked “gene”) and in the UMAP data visualization ^16^ (Figure 1G).

Because the profiles of most variants tend to cluster together within each gene, as observed in the hierarchical clustering of the correlation matrix (Figure 1F), we conclude that the phenotypic variations of alleles remain closely related to the reference gene and rarely result in a major phenotypic disruption that places them in a different cluster. This type of closely related variation is consistent with previous studies in morphological and transcriptional profiling ^14,17^, which report that the major factor of variation detected by profiling platforms is first associated with cell lines, then with groups of perturbations that share similar mechanisms, and finally with specific effects of each perturbation.

Interestingly, for a subset of alleles with functional annotation, Cell Painting profiles cluster the data in two major parts in the correlation matrix (Supplementary Figure 1): one part is enriched with variants from known oncogenes such as BRAF, EGFR, KRAS, and CTNNB1, and the other part is enriched with variants from known tumor suppressor genes, including FBXW7, KEAP1, and STK11. This result confirms that morphology captures relevant cellular changes associated with known cancer biology.

### 3. Cell morphology-based variant impact phenotyping (cmVIP) correctly classifies benchmark alleles

We next tested whether the detected differences in morphology can predict each variant’s impact on gene function. Using the decision tree from prior expression-based variant impact profiling (eVIP) ^6,7^, we tested for significant differences in the similarity between cell morphological profiles of reference genes and their variants (Methods). We call this extension of VIP *cell morphology-based VIP* (cmVIP), which interprets replicate correlations among alleles as probability distributions that can be compared using statistical tests (Figure 2A, Methods).

**Figure 2.**
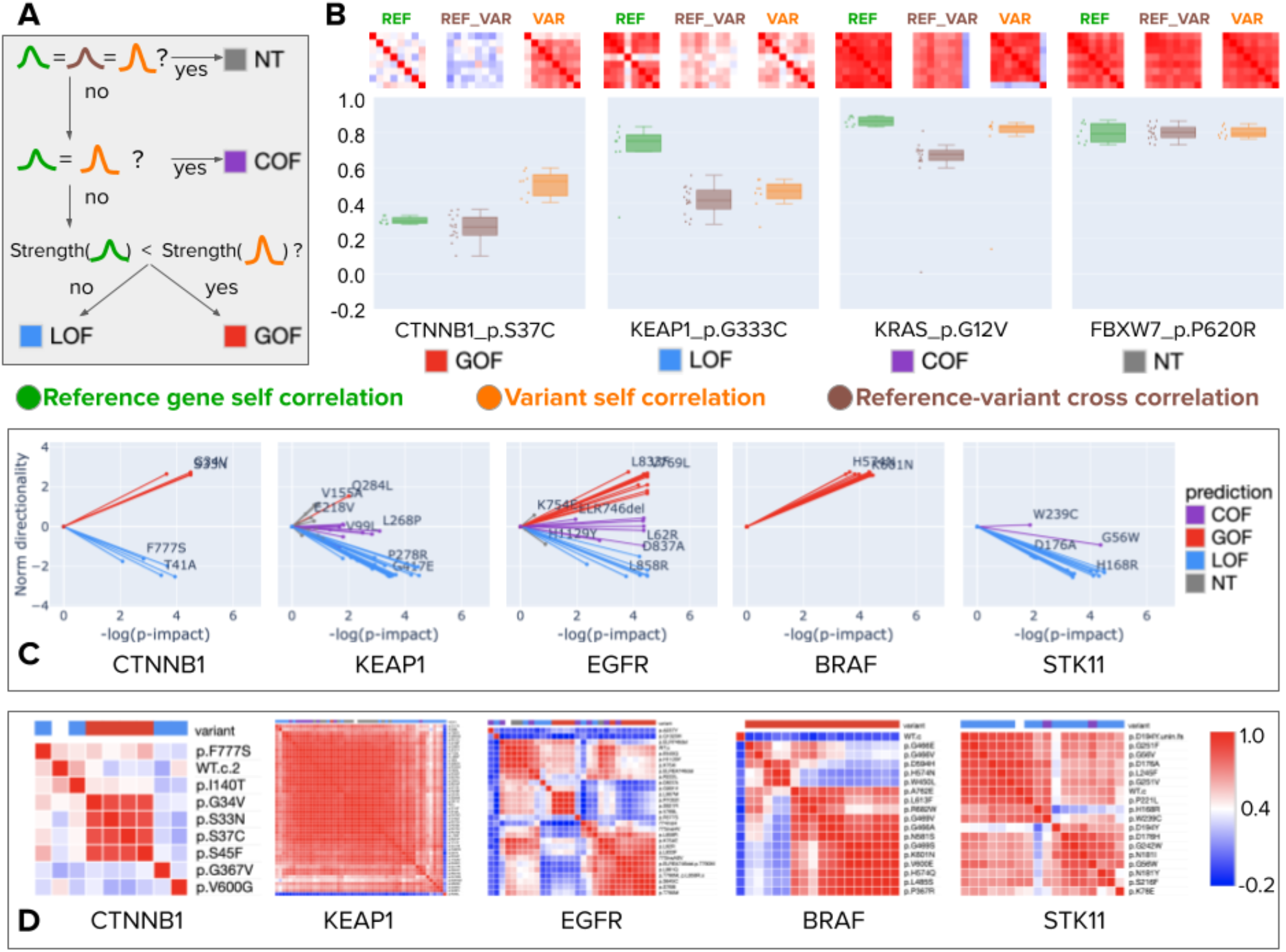
Morphology-based variant impact phenotyping (cmVIP) and resulting predictions in a diverse set of genes and alleles. A) Decision tree of the VIP algorithm ^6,7^, which we adopt for classifying variants by their Cell Painting profiles as gain of function (GOF), loss of function (LOF), change of function (COF) and neutral (NT) mutations. B) Example predictions by cmVIP on four alleles, one of each type. The correlation matrices at the top show how similar the replicates of each pair are (reference gene self-correlation, reference-variant cross-correlation, and variant self-correlation). The correlation matrix colors represent the correlation values in the same color scale as in D. The box plots below the matrices show the distribution of median values of the matrices’ rows (self-correlation) and columns (cross-correlation). C) Sparkler plots display the magnitude and directionality of predictions for all alleles in a gene set. The x-axis represents the negative log p-value of the impact test (the larger the more impactful), and the y-axis represents the log p-value of the directionality test polarized by the result of the strength test. All alleles for these genes are displayed, but only a few are annotated to aid visualization. All the plots and annotations can be queried at full scale in the interactive website: http://broad.io/cmvipD) Correlation matrices for the groups of alleles presented in C.

We found that cmVIP correctly classified 100% of the set of 20 well-characterized alleles (Supplementary Table 1) that Berger et al. previously used in evaluating eVIP. This set of 20 alleles has been previously characterized using functional assays. We also predicted the directionality of the alleles in this benchmark set and found that cmVIP correctly classifies 16 out of the 20 alleles in one of two groups: change of function (COF) or gain of function (GOF) variant vs loss of function (LOF) variant (Supplementary Table 1).

Finally, we also estimated the false positive rate of cmVIP with mock alleles using a set of high-replicate controls. We collected 64 replicates for each of these control alleles (known to have high phenotypic activity), and then we sampled random groups of 8 replicates without replacement to simulate reference genes and variant pairs. Next, we run the cmVIP analysis to determine if this mock pair has an impact, and we expected a negative answer as a result. We ran this simulation 1,000 times and found that cmVIP falsely calls the mock alleles impactful 6.75% of the time on average (Supplementary Table 2), close to the false discovery rate of 5% at which the testing procedure is controlled. These results suggest that Cell Painting can reliably predict the impact status of variants of unknown significance.

### 4. The impact of variants of unknown significance can be predicted at high-throughput with Cell Painting

We next explored the full set of genes and alleles. cmVIP found 258 alleles (79.3%) to be impactful; from these 158 alleles (48.6%) were classified as GOF or COF variants, and 100 as LOF variants (30.7%). We show examples (Figure 2B) and provide an online resource to explore all genes and their variants (http://broad.io/cmvip).

Similar to eVIP ^6,7^, the cmVIP decision tree (Figure 2A) starts by looking at the correlation matrices of reference gene replicates (REF self-correlation) and variant replicates (VAR self-correlation) as probability distributions. Given that the image-based profiling workflow involves control-based normalization (Methods), we expect self-correlation matrices (correlation values between true replicates) to have high signal when the underlying phenotype is different from negative controls. This interpretation applies to reference gene and variant self-correlation matrices (REF_REF and VAR_VAR in Figure 2B). Finally, the reference gene vs variant cross-correlation matrix (REF_VAR) reveals how similar is the variant in question to its corresponding reference gene.

cmVIP interprets statistically significant changes in these three distributions of similarities among replicates in a biologically meaningful way. For instance, CTNNB1 has a relatively low signal in its reference form (REF_REF median signal strength = 0.30, Figure 2B), meaning overexpressing it in cells changes their morphology only marginally. A gain-of-function (GOF) variant in this gene (e.g. CTNNB1 S37C in Figure 2B), by contrast, yields a relatively stronger signal (VAR_VAR median signal strength = 0.50, Figure 2B) and is different from the reference (REF_VAR median signal strength = 0.25). Loss-of-function (LOF) variants, on the other hand, are usually characterized by alleles with a weak phenotype, relative to a reference that has a strong phenotype (e.g. KEAP1 G333C, Figure 2B). Change-of-function variants show strong phenotypes for the reference gene and variants, and they differ from each other (e.g. KRAS G12V, Figure 2B). Finally, neutral mutations show high similarity between the reference gene and variant, indicating no detectable phenotypic change (e.g. FBXW7 P620R, Figure 2B).

The statistical tests of cmVIP provide p-values for such differences, which can be visualized to compare the impact and directionality of a group of variants using sparkler plots (Figure 2C). These show, for example, that the KEAP1 and STK11 alleles tested in our study mainly present a LOF or COF variant pattern; BRAF alleles have a GOF behavior, while CTNNB1 and EGFR alleles present a diverse range of GOF, COF, and LOF variants.

### 5. Cell Painting reveals allele heterogeneity at single-cell resolution

Image-based profiling inherently offers single-cell resolution while being the lowest cost even among bulk profiling methods. We investigated whether single-cell morphological profiling might provide insights into the heterogeneity of allele subpopulations or other phenotypic mechanisms that cannot be observed using bulk-level data ^18^. We extract single cells from Cell Painting images using the seeded-watershed segmentation algorithm and then compute deep learning feature embeddings for each one individually (Methods). The feature representation of single cells has been transformed using a spherizing transformation with respect to a set of 1.5 million negative control cells to minimize the impact of technical variation across batches.

We found that single-cell data visualizations for each allele allow qualitatively observing cell heterogeneity and the relationship among cells overexpressing a particular allele relative to its reference gene counterpart. For example, the two BRAF alleles in Figure 3 (V600E and W450L) were classified as impactful GOF variants using the cmVIP algorithm: both showed a strong phenotype in the variant replicate correlation matrices compared to the reference gene, whose replicate correlation was weak (Figure 3A,D). When looking at single cells in reduced-dimensional space (Figure 3B,C), we observe that each alleles’ phenotypes move to different regions of the phenotypic space compared to the reference gene. These two different regions are not exclusive of these two alleles; they are also occupied by other BRAF variants (W450L is similar to H574N and D594H, while V600E is similar to L485S, K601N, and H574Q; interactive website http://broad.io/cmvip/variants/BRAF_p.W450L/). This suggests different mechanisms between the two groups of alleles; in fact, it is well-known that V600E and other constitutively activating alleles have different behavior than W450L and other variants of the same gene ^19–21^.

**Figure 3.**
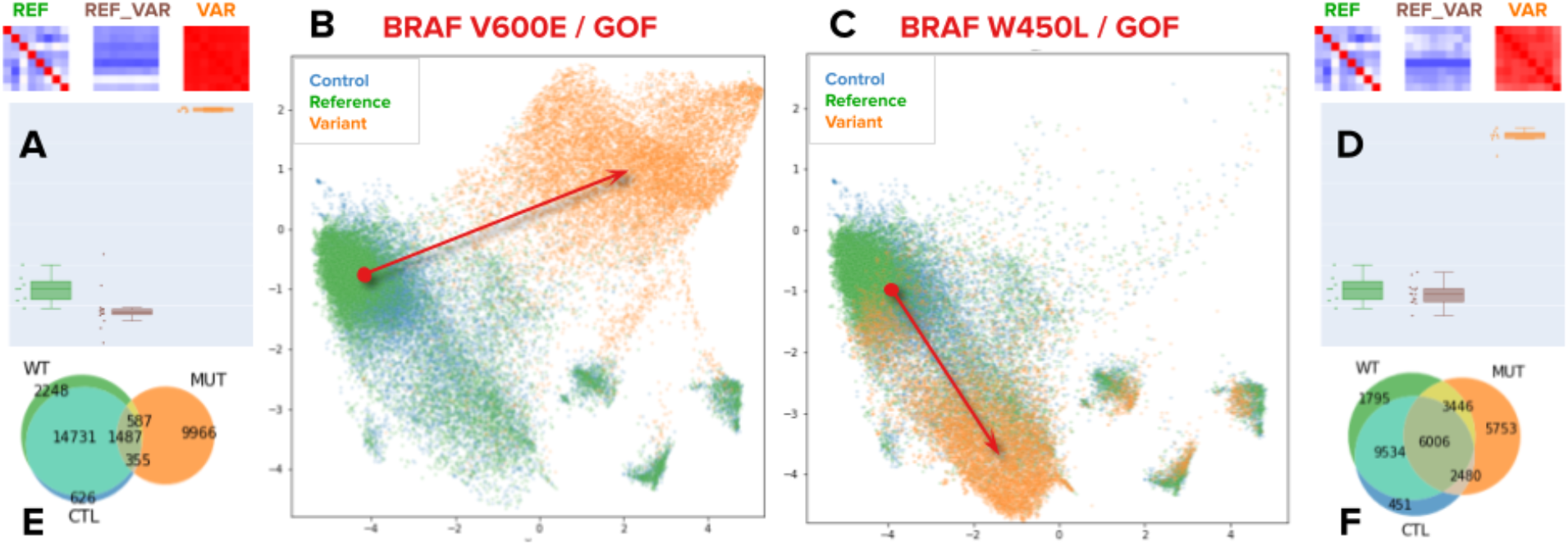
Single-cell heterogeneity of variants. Different mutations of the same gene result in different phenotypes. A,D) Correlation matrices and box/dot plots of bulk-level profiles for the corresponding alleles, as in Figure 2. These matrices are used to obtain the impact and directionality predictions with cmVIP. B,C) UMAP visualizations of three populations of cells, the empty control population (in blue), the reference gene population (in green), and the variant population (in orange). Each point in the plots is a single cell extracted from the Cell Painting images using segmentation. The UMAP embedding for all panels is computed using a fixed sampling of negative control wells. Arrows indicate the shift in phenotypic space from the reference gene population to the variant population. Note that variants of the same gene move in different directions. E,F) Venn diagrams of the overlap among the reference gene, variant and control populations of cells. These counts are obtained using graph analysis in the original feature space (Methods).

We quantify and summarize these variations in single-cell states using graph analysis and nearest neighbors (Methods), which can be observed in the Venn diagrams (Figure 4E,F) that summarize single-cell counts that have shared phenotypes (Methods). UMAP plots that allow single cell visualizations, as well as the corresponding Venn diagrams are available for all the variants in our study at http://broad.io/cmvip.

**Figure 4.**
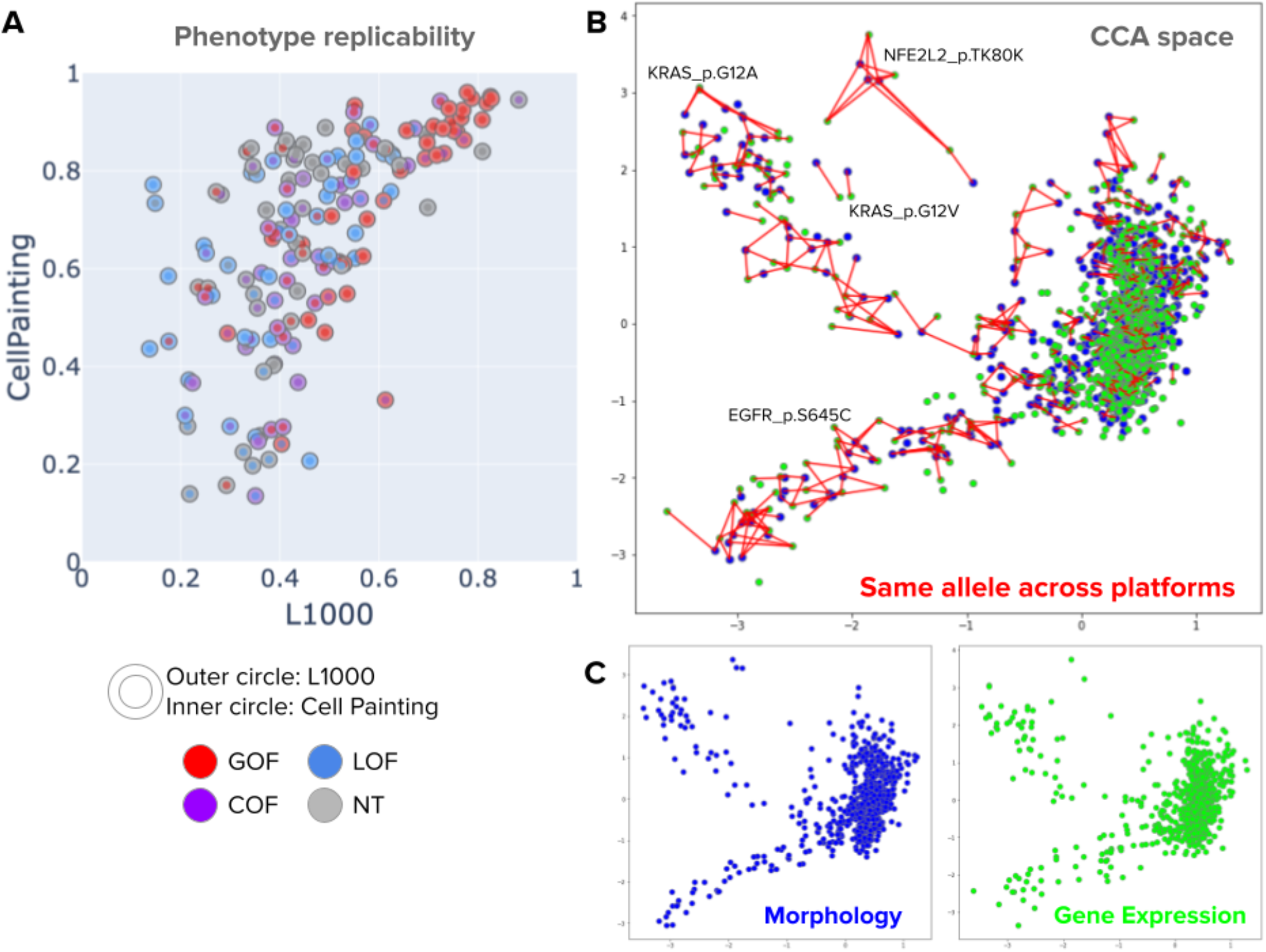
Correlation between Cell Painting profiles and L1000 profiles for a common subset of 160 variants. A) Signal replicability, defined as the median pairwise correlation between replicates of the same allele, was calculated for each variant in the common subset in both profiling platforms. The x-axis corresponds to the signal strength in L1000 and the y-axis represents the signal strength in Cell Painting. The Spearman correlation coefficient is 0.69. B) Canonical correlation analysis (CCA) in the multidimensional feature space for both profiling platforms at the perturbation level. CCA obtains a common latent space by finding the directions of maximal correlation between two multivariate datasets, allowing us to project data points from Cell Painting and L1000 in the same subspace. The axes in this plot are the first and second CCA directions. Points in blue are morphology profiles and points in green are gene expression profiles. The red lines connect two points of different modalities that represent the same gene or allele. C) Same representation of Cell Painting profiles (morphology) and L1000 profiles (gene expression) in CCA space as in B, but using independent plots for each platform.

### 6. Cell Painting phenotypic variations are highly correlated with gene expression variations

A subset of 160 alleles that we profiled for this study was previously profiled using transcriptional profiling with the L1000 platform. Given the pairs of profiles for the same perturbations, we investigated the extent to which phenotypic variation captured with Cell Painting profiles corresponds with L1000 variation. Although they are not identical, we found high correlation between both platforms in this subset of alleles by conducting two different correlation analyses (Figure 4).

First, when measuring the phenotype replicability of alleles, we found a high correlation between the signal of Cell Painting profiles and the signal of L1000 profiles (Figure 4A). Phenotype replicability is defined as the median replicate correlation among true replicates of the same allele; high correlation values indicate that the underlying condition is detectable by the profiling platforms and reproducible among replicates, i.e. when an allele has a high signal in L1000 it is likely to be detected with high signal in Cell Painting as well.

Second, we projected perturbation-level profiles of both platforms to the same latent space using canonical correlation analysis (CCA), which finds directions of maximal correlation between two paired multidimensional datasets. We found high agreement between profiles from both platforms when projected into the first two CCA components (Figure 4B,C). This alignment confirms that the relative similarities and differences observed between allele phenotypes in our study can be reproduced with different assays under different experimental settings, increasing the confidence that the signal captured by both platforms is reliable and biologically meaningful.

### 7. Cell Painting predictions are consistent with transcriptional profiling predictions

We next explored how well cmVIP’s predictions matched known observations about cancer genes and alleles. Beyond the 20 benchmark genes tested above (Supplementary Table 1), 140 additional alleles in our study were previously characterized using transcriptional profiling via expression-based variant impact phenotyping (eVIP) ^6^.

We found that eVIP and cmVIP platforms agree on the predicted impact for 123 of the 160 alleles (76.8%) (Figure 5A). From those concordant predictions, 102 alleles were found to be impactful while 21 were found to be neutral. This level of agreement increases the confidence that both phenotypic profiling platforms are consistently quantifying relevant cancer biology in the underlying experiment, and also confirm that the VIP strategy generalizes well to diverse phenotypic readouts.

**Figure 5.**
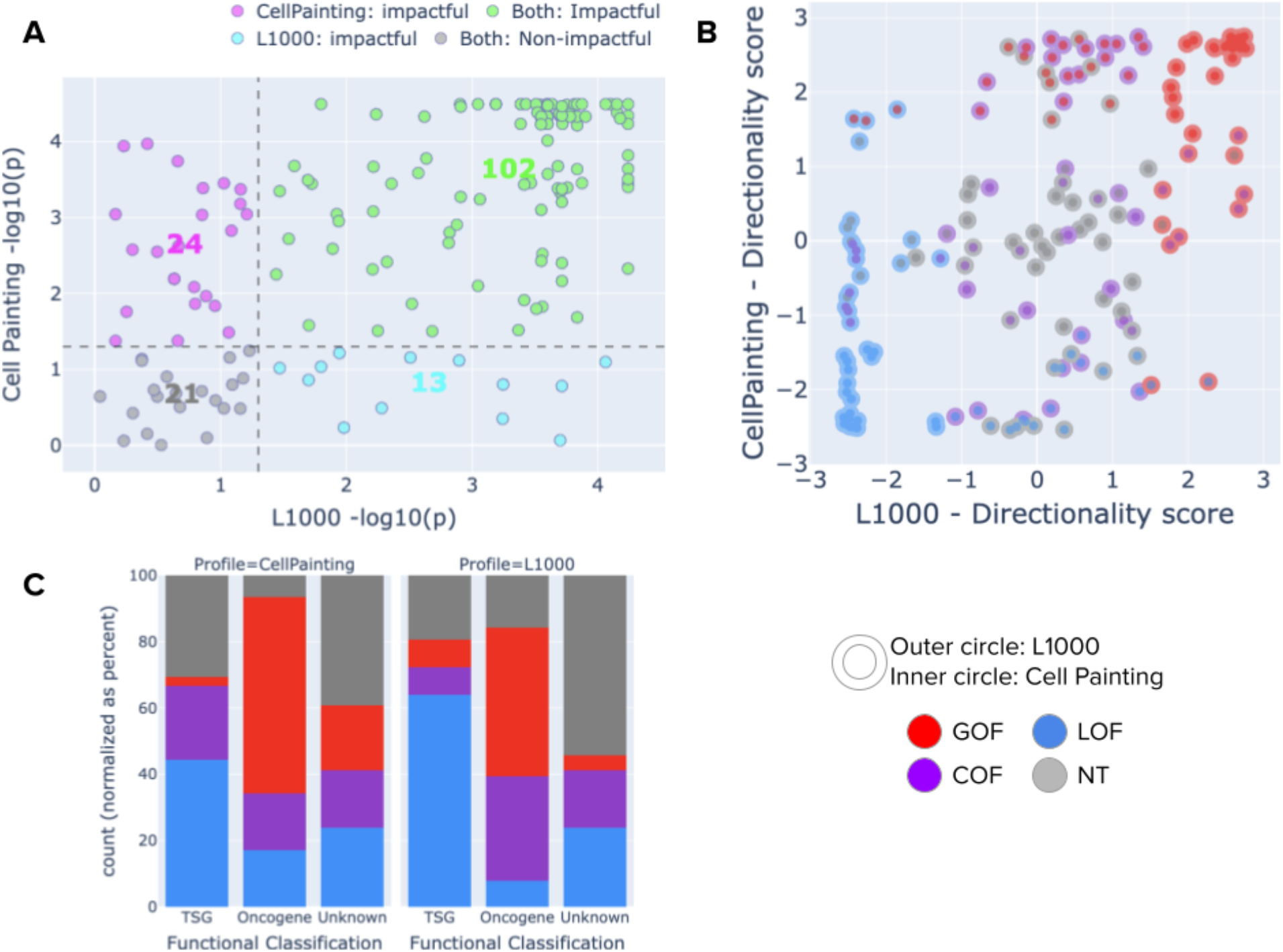
Comparison of VIP predictions using Cell Painting (morphological profiling) and L1000 (transcriptional profiling). Both platforms use the same underlying statistical tests of the VIP algorithm. A) Impact test results. The x-axis presents the negative log p-value obtained by eVIP (L1000), and the y axis represents the negative log p-value obtained by cmVIP. The dotted lines represent the significant threshold considered in this study (0.05). Each point is one allele and its color indicates the prediction agreement between the two platforms: green is impactful by both platforms, gray is neutral by both platforms, pink is impactful by Cell Painting only, and blue is impactful by L1000 only. B) Directionality test results. The x-axis indicates the polarized log p-value obtained by L1000, and the same for Cell Painting in the y axis. Each point is one allele with the inner circle colored according to the predictions obtained by each platform. C) Distribution of cmVIP and eVIP predictions in known oncogenes, known tumor suppressor genes (TSGs), or genes of unknown function. The distribution of oncogenes is enriched with GOF/COF calls in both platforms, and similarly, the distribution of tumor suppressor genes is enriched with LOF calls.

Next, we evaluated the agreement between both platforms in the predicted directionality of impactful alleles, and we found consistency in 21 LOF variants, 29 GOF variants, and 7 COF variants (Figure 4B). A common disagreement appears with alleles that are called GOF by one platform and COF by the other (23 variants). Other disagreements are observed between LOF vs NT (17 variants), and COF vs NT (10 variants), which happen when one platform has higher phenotype strength for those alleles than the other, i.e., one platform detects the phenotype and the other does not. A few unexpected disagreements also appeared in 5 cases with LOF vs GOF directionality classifications: IDH2 K130del (CP:GOF / L1000:LOF), IDH2 S249G (CP:GOF / L1000:LOF), PIK3CA E600K (CP:GOF / L1000:LOF), RIT1 R122L (CP:LOF / L1000:GOF), and CTNNB1 V600G (CP:LOF / L1000:GOF). These may either represent occasional technical errors, or cases where the function of the WT or variants allele is undetectable by one platform versus the other.

Finally, we looked at the functional classification of genes for a few alleles in the common set (Figure 5C). Our set of 160 alleles in common between both platforms has not been completely characterized as to their GOF, LOF, COF, NT status, but many of their genes are classified as tumor suppressors or oncogenes. One would expect that alleles found in tumor suppressors are more likely to be LOF than GOF/COF whereas alleles found in oncogenes are more likely to be GOF/COF than LOF. We found that both cmVIP and eVIP make predictions consistent with these expected trends (Figure 5C).

## Discussion

Here we demonstrate that images of cells overexpressing given cancer-associated variants can be used to predict their impact on a diverse array of genes’ functions at high-throughput using the cmVIP strategy. The signal obtained from image-based profiling was sensitive to morphological variations of lung cancer variants in this experiment and was useful to characterize and make predictions for 325 alleles. The accuracy appears comparable to that of transcriptional profiling, and the two platforms’ predictions are generally concordant. Resolving the impact of variants at high throughput has the potential to accelerate precision oncology ^22,23^.

Unbiased cell morphological profiling based on the Cell Painting assay has been shown to be a powerful approach for drug discovery and functional genomics ^10,24^. Our work expands the application of image-based profiling with Cell Painting to cancer variant phenotyping, indicating that it might be scaled up to much larger collections of variants efficiently and cost-effectively. The approach may be extended from somatic variations found in cancer to investigate the impact of germ-line variations of unknown significance in humans. Exploring a variety of cell lines and examining their concordance for variant impact prediction would be particularly interesting.

Image-based profiling provides single-cell resolution to investigate cellular heterogeneity across perturbations. We observed single-cell phenotypic differences between variants of the same gene, which could provide insights into functional differences of alleles. The richness of single-cell variation and the ease of implementation suggests that phenotypic studies could be performed using image-based profiling with fewer technical replicates while maintaining the ability to detect meaningful morphological variations. We leave it to future research to further investigate particular cases where single cells reveal interesting heterogeneity patterns to uncover novel cancer biology, as well as potential confounders therein.

In this work, we also used novel computational methods based on deep learning models to transform images of cells into quantitative phenotypic profiles, an approach just starting to be used in the field ^25^. The sensitivity of image-based profiling can be further increased with the advent of more powerful machine learning algorithms that extract precise patterns from images using computer vision. Our methods are open source and can be adopted for similar applications in the future, and we also expect contributions from the imaging community to develop new techniques that harness the morphology of cells for studying cellular biology.

Future studies might aim to integrate imaging and mRNA data types (if both are available) to explore whether their predictive power increases when combined. Our results indicate that morphology and gene expression, as captured by the Cell Painting and L1000 assays, measure highly correlated phenotypic variation, which mutually confirms their ability to detect meaningful biological events. This suggests the possibility to model their correspondences using computational approaches to translate one data type from the other or to understand their causal relationships. Our dataset has been simultaneously used in a study to identify which gene expression variations correspond with which morphology variations, and vice versa ^26^. While this has been explored at the bulk level, our results and previous work based on scRNAseq ^18^ indicate that this type of analysis could be extended to understand multi-omics connections at the single-cell level.

We publicly provide all data used and created in this study, including the raw images and the computed profiles (Methods). Further, we provide a public portal where researchers can explore genes and alleles of interest to see the distribution of signal strength, impact and directionality predictions, VIP calls, and UMAP plots of genes and alleles (http://broad.io/cmvip/).

## Acknowledgments

We thank Mukta Bagul, JT Neal, and Oana Ursu for helpful discussions. Funding for the project was provided by the National Institutes of Health (NIH R35 GM122547 to AEC), the Broad Institute V2F Initiative (to AEC), the Broad Institute Schmidt Fellowship program (to JCC), and the Slim Initiative for Genomic Medicine, a project funded by the Carlos Slim Foundation in Mexico. The authors declare no competing financial interests.

## Methods

### Profiling cancer variants with Cell Painting

Cells were grown, stained, fixed, and imaged as described in our protocol ^11^. Briefly, A549 cells are grown in 384-well format and infected with lentiviral ORF constructs which induce over-expression of various ORFs and alleles therein. After 96 hours, MitoTracker stain was added to live cells to label the mitochondria. Cells were then fixed, fixed with formaldehyde, permeabilized with Triton X-100, and stained with the remaining dyes to identify the nucleus (Hoechst), nucleoli and cytoplasmic RNA (SYTO 14), endoplasmic reticulum (concanavalin A), Golgi and plasma membrane (wheat germ agglutinin), and the actin cytoskeleton (phalloidin). Plates were imaged using an ImageXpress Micro XLS automated microscope (Molecular Devices). We captured images from nine fields of view (sites) per well in five fluorescent channels each using a 20× lens. Separate, grayscale image files for each channel were then stored in 16-bit TIFF format. All raw image data are publicly available at the Cell Painting Image Collection (https://registry.opendata.aws/cell-painting-image-collection/).

The alleles in the ORF library represent a subset of those identified in an analysis of 412 primary lung adenocarcinomas that were previously sequenced 9,25, which detected 518 unique missense and in-frame insertions or deletions in the 50 genes prioritized in this study 26. In all, ORF constructs for 325 variants (and reference versions) of these 50 genes were successfully generated and assayed. An additional 88 constructs are included in the dataset, representing TP53 alleles that inadvertently had double mutations. A comprehensive description of the process for selecting the constructs that were analyzed is presented in Supplementary Figure 2. The additional alleles have been included in the dataset for completeness. Eight replicates were assayed for two of the plates of constructs; a third plate–comprising multiple replicate wells of a small number of “control” alleles–was assayed in two replicates.

### Cell line

A549 cells (adenocarcinomic human alveolar basal epithelial cells), RRID:CVCL_0023, were obtained from ATCC; they were not additionally authenticated prior to this experiment. The cell line tested negative for mycoplasma prior to this experiment.

### Mutated cDNA Library

The cDNA library is identical to that described in ^6^: wild-type ORF constructs were obtained from the human ORFeome library version 5.1 (http://horfdb.dfci.harvard.edu) and used as templates for site-directed mutagenesis to generate mutated cDNAs in the pDONR223 Gateway entry vector. All constructs used in downstream analyses were validated by Sanger sequencing to include the intended mutation and no other identified sequence differences relative to the wild-type construct. After sequence verification, mutated ORFs were shuttled into the pLX317 lentiviral expression vector by LR recombination.

### Image analysis

#### Illumination correction

TIFF images were corrected for non-homogeneous illumination variation across the image field using a retrospective approach ^27^. Briefly, the method computes illumination correction functions by averaging all images of the same channel in a multi-well plate, followed by a median filter. Images in the plate are corrected by dividing their intensity values by the corresponding illumination correction function. For visualization purposes (e.g. example images reported in Figure 1), we rescale intensity values to fit the range of 255 grayscale values separately for each channel.

#### Segmentation

Single cell identification was performed using CellProfiler ^28,29^ with the Identify Primary (nuclei) and Secondary (cell bodies) objects functionality. This approach runs thresholding and seeded watershed to identify the structures of interest. The single cell analysis presented in this work was conducted by recording the center of the nucleus of each cell and then cropping a fixed-size region around these coordinates (see feature extraction below). Cell masks were not used to isolate cells from the background.

#### Feature extraction

Feature extraction computes a numerical representation of the image content. Standard approaches use handcrafted descriptors such as texture or shape features ^28^. Although widely used to quantify cellular morphology, they still require careful hyperparameter tuning to get high-quality representations, and, due to the high variability in the acquisition process, different datasets require custom adjustment. In contrast, representation and deep learning methods aim to automatically find transformations that yield a compact and meaningful representation based solely on image pixels. Previous empirical exploration showed promising results using deep learning models trained in the natural images and then using them to extract features from cellular images ^12^. Motivated by this and the success of transfer learning in computer vision applications, we use a pre-trained EfficientNet neural network ^30^ to obtain embeddings for Cell Painting images.

First, we compute a feature vector that represents the content for each segmented cell. Bounding boxes are centered around the center of segmented cells and cropped to 128×128 pixels, and re-scaled to 224×224 pixels to match the expected input of EfficientNet B0. We process each of the five Cell Painting channels independently as if they were separate RGB images by replicating their grayscale values in three channels and then running them through the EfficientNet. We keep the feature vectors of the second-to-last layer, which produces a 1,280 dimensional representation for one image, and then concatenate the five vectors (one per channel), generating 6,400 features to represent a single-cell profile. This process was executed using the DeepProfiler open source tool (https://github.com/cytomining/DeepProfiler).

### Image-based profiling

In general, we followed the image-based profiling best practices defined by the community for transforming images into quantitative readouts ^31^. More specifically, in order to get perturbation-level (or bulk-level) profiles, we first aggregate single-cell profiles into replicate-level (or well-level) profiles by computing their mean, and then aggregate replicate-level profiles by computing their median. In our study, we conducted a multi-level analysis of image-based profiles including perturbation-level profiles to verify associations among alleles and with gene expression data; replicate-level profiles to make impact and directionality predictions using the cmVIP algorithm; and single-cell level profiles to explore phenotype heterogeneity.

### Data normalization and batch correction

As is the case in many biological experiments, imaging assays may also be prone to nuisance variation due to technical artifacts. We used *negative control spherizing* to correct for batch effect biases, which has shown to be effective in other studies ^13,14,32^. The spherizing transform used in this work makes the assumption that negative controls sampled from different batches ought to be similar to each other in the biological sense, and any deviations from this normal looking phenotype is rather technical. Therefore, by finding a new embedding space where controls have roughly the same amount of variation in every dimension, the patterns of interest naturally emerge while batch effects are minimized. This is the same principle used in the Typical Variation Normalization (TVN) transform ^13^.

Spherizing is achieved by computing a singular value decomposition of the covariance matrix of control profiles and then scaling all the directions of the orthogonal basis by the inverse of the corresponding eigenvalues ^33^. The rescaled dimensions define a new representation space where large variations (usually associated with nuisance variations) are reduced, and rare variations (usually phenotypic variations) are amplified. We calculated the transformation matrix using control samples at the replicate-level, and used it to project all other perturbation profiles in our experiment into the corrected feature space. The spherizing transform has a regularization parameter for safely inverting the eigenvalues of the covariance matrix, which was set to 0.01 in our analysis.

### Cell morphology-based Variant Impact Phenotyping (cmVIP)

Our procedure closely follows the eVIP algorithm ^6,7^. For any given variant and its corresponding reference gene, cmVIP estimates the impact and directionality of the variant based on three correlation sets: 1) variant self-correlation: median correlation values in the rows of the replicate correlation matrix of the variant, 2) reference gene self-correlation: median correlation values in the rows of the replicate correlation matrix of the reference gene, and 3) reference-variant cross correlation: median correlation values in the rows and columns of the correlation matrix between variant and reference gene replicates.

cmVIP follows the rule-based decision tree depicted in Figure 2A. The first stage determines if there is a statistically significant difference between any of the three correlation sets using the Kruskal-Wallis test, which is a non-parametric test. If the test rejects the null hypothesis, i.e. there is a difference, then the variant is considered to be impactful, otherwise, the variant is considered to be neutral.

For impactful variants, cmVIP determines its functional directionality by running a Wilcoxon statistical test on variant self-correlations vs reference gene self-correlations. If the test rejects the null hypothesis, i.e. there is a difference between variant and reference gene, then their medians are directly compared. If the median of the variant is higher than the reference one, we predict it is a gain-of-function variant, otherwise, we call it a loss-of-function variant. In case that the Wilcoxon test fails to reject the null hypothesis, i.e. there is no difference between variant and reference, we predict it is a change-of-function variant.

The Benjamini-Hochberg multiple-hypothesis correction procedure is used to control the false discovery rate (FDR) of each step to be less than 5%.

### Single-cell analysis

We used single-cell profiles to explore phenotypic differences between variants of the same reference gene. The first step before using single-cell profiles for quantitative analysis was to spherize the control distribution at the single-cell level (see Data normalization and batch correction for more details). To accomplish this, we used approximately 1.5 million single-cell profiles taken from all the 320 control wells in our experiment to compute the spherizing transform. Then, we projected all other single cells coming from overexpression perturbations in the corrected space. The regularization parameter used for spherizing single cells was set to 0.01 (same as in the aggregated profiles case).

Corrected single-cell profiles were then used to compute visualizations using the UMAP projection one gene at a time, including the reference gene and all its available variants. We observed that, when coloring single cells in this UMAP visualization with plate identifiers, the different replicates are well mixed and integrated (random coloring patterns, see http://broad.io/cmvip for examples). By computing visualizations for all alleles of the same gene at the same time, we can also qualitatively assess the relative differences among their phenotypes. We used the UMAP algorithm default parameters in their Python implementation in all cases to reveal the structure of the feature space in the most unbiased way possible.

Beyond qualitative single-cell analysis using UMAP visualizations, we used graph analysis based on nearest neighbors to objectively quantify the overlap between populations of cells in the original feature space. In this analysis, we first created a five-nearest neighbor graph using a sample of 15,000 single cells coming from three populations (5,000 from each): reference gene, variant and negative controls. The sample from each population comes from a mix of all replicates. In this graph, we proceed to classify the phenotype of single cells in one of seven categories: 1) pure reference gene phenotype, 2) pure variant phenotype, or 3) pure control phenotype, if all the five nearest neighbors are from one of these three populations. 4) Shared reference-variant phenotype, 5) shared reference-control phenotype, or 6) shared variant-control phenotype, if the five nearest neighbors are a mix of these two populations. Finally, 7) combined phenotype, if the five nearest neighbors are a mix of the three populations. The classification of single cells in these seven categories is used to create the Venn diagrams of single cell phenotypic overlap presented in Figure 3 and in the interactive website http://broad.io/cmvip.

## Data and code availability

We make the data used in this project publicly available. The raw images can be downloaded from the AWS Open Data - Cell Painting Image Collection (https://registry.opendata.aws/cell-painting-image-collection/ in the following path: cytodata/datasets/LUAD-BBBC043-Caicedo/). CellProfiler was used to prepare and segment cells. The code used to process raw images and obtain deep learning features, which is based on TensorFlow ^34^, is available at https://github.com/cytomining/DeepProfiler/.

After obtaining image-based profiles, all our analysis was developed using the data science Python ecosystem, including NumPy ^35^, SciPy ^36^, Pandas, and JupyterLab, among others. All our scripts and notebooks are available at https://github.com/broadinstitute/luad-cell-painting. Finally, an interactive website with the aggregated data, predictions for all alleles, and full resolution figures presented in this manuscript is available at http://broad.io/cmvip.

## Supplementary material

**Supplementary Figure 1.**
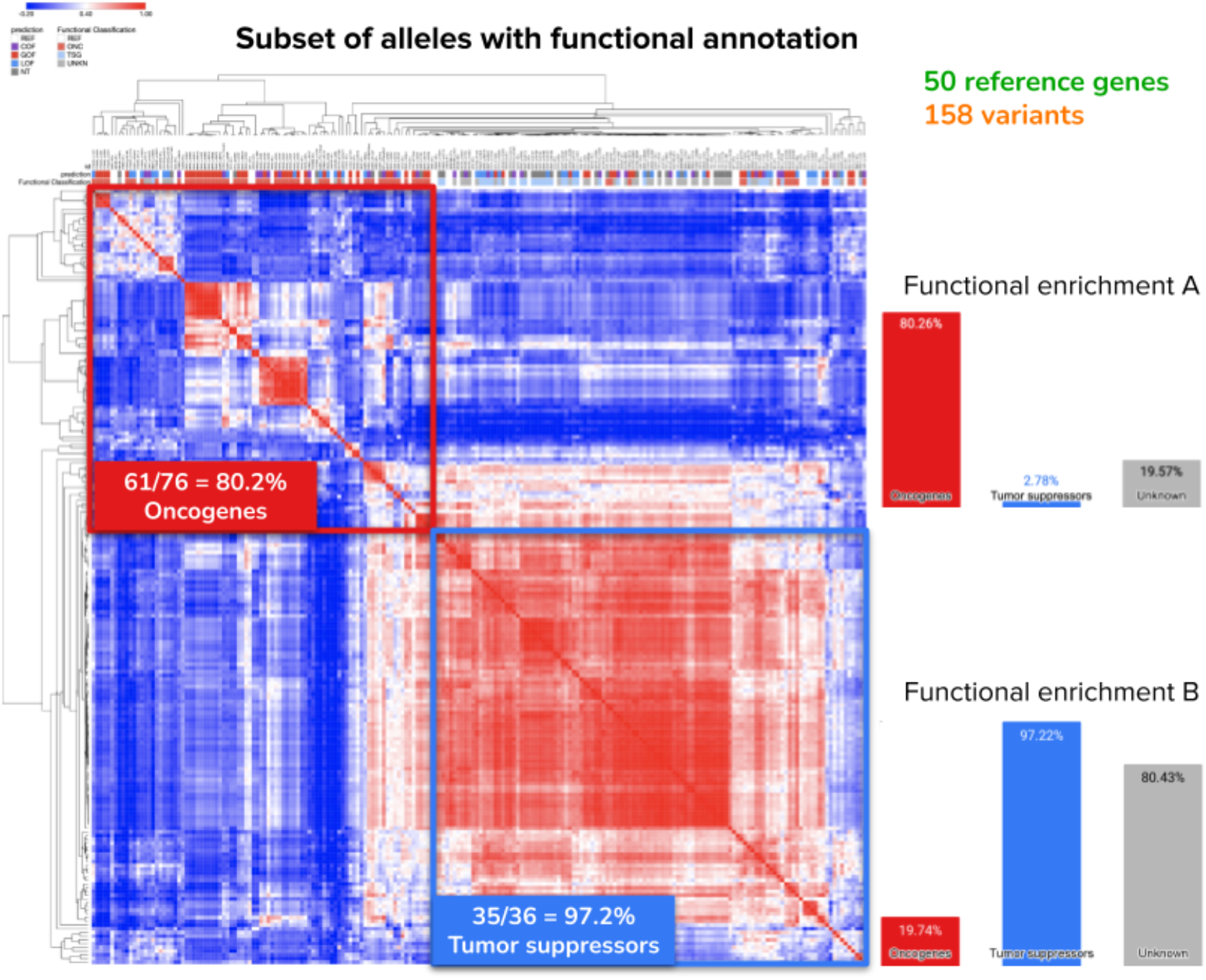
Correlation matrix of Cell Painting profiles for a subset of alleles with functional annotation. After ordering rows and columns according to the hierarchical clustering, the matrix can be divided in two parts: one enriched with oncogenes and the second enriched with tumor suppressor genes. Enrichment here is defined as the proportion of alleles in one functional category that are present in the group with respect to all alleles of that category.

**Supplementary Figure 2.**
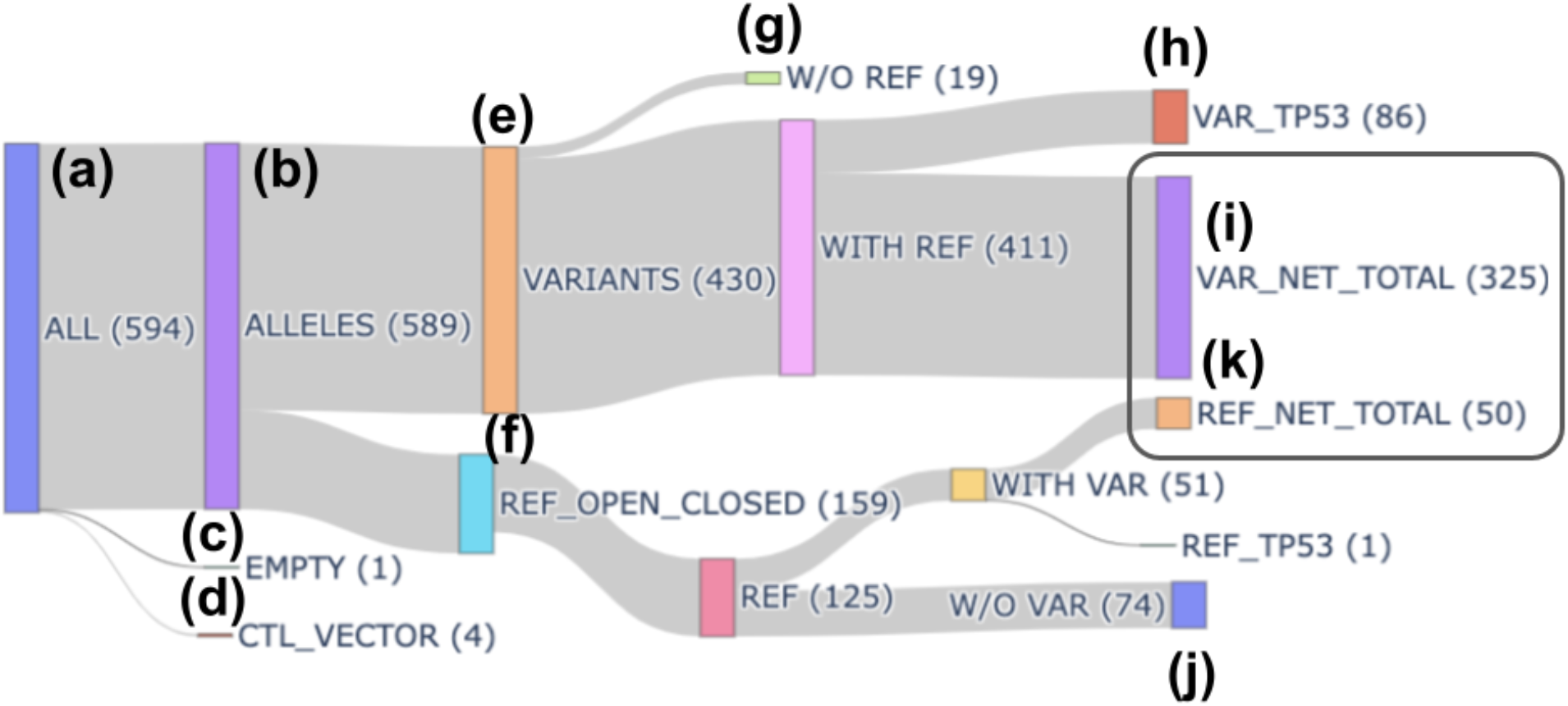
Disaggregation of the metadata in the raw dataset. The *x_mutation_status* column tags the content of each assay. It contains 594 unique values (a), 589 (b) correspond to alleles, EMPTY (c) corresponds to the control tag, and ctl_vectors (d) are not considered in this analysis. Out of the 589, 430 (e) are VARIANTS and 159 (f) are REFERENCE alleles. We discarded 19 variants (g) without reference, and 86 TP53 Variants (h) resulting in the 325 alleles(i). We also discarded 74 references that do not match with any variant (j), ending up with 50 (k) REFERENCE alleles. The enclosed subset represents the data used in this study.

**Supplementary Table 1.** Benchmark classification results.

| Allele | Reference | L1000 | CellP | Prior annotation | Allele | Reference | L1000 | CellP | Prior annotation |
| --- | --- | --- | --- | --- | --- | --- | --- | --- | --- |
| ARAF_p.S214F | Imielinski et al., 2014 | COF | LOF | GOF | KRAS_p.G13V | Prior et al., 2012 | GOF | COF | GOF |
| ARAF_p.S214C | Imielinski et al., 2014 | COF | COF | GOF | KRAS_p.G12R | Prior et al., 2012 | GOF | GOF | GOF |
| ARAF_p.D429A | Imielinski et al., 2014 | GOF | COF | LOF | KRAS_p.G12S | Prior et al., 2012 | GOF | GOF | GOF |
| CTNNB1_p.S37C | Palacios et al., 1998 | GOF | GOF | GOF | KRAS_p.G12D | Prior et al., 2012 | GOF | COF | GOF |
| EGFR_p.ELREA746del | Greulich et al., 2005 | COF | COF | GOF | KRAS_p.G12F | Prior et al., 2012 | GOF | GOF | GOF |
| EGFR_p.L858R | Greulich et al., 2005 | NT | LOF | GOF | NRAS_p.Q61L | Prior et al., 2012 | COF | GOF | GOF |
| KEAP1_p.G333C | Hast et al., 2014 | LOF | LOF | LOF | RIT1_p.F82L | Berger et al., 2014 | COF | COF | GOF |
| KRAS_p.G12Y | Prior et al., 2012 | COF | GOF | GOF | RIT1_p.R122L | Berger et al., 2014 | GOF | LOF | GOF |
| KRAS_p.G12A | Prior et al., 2012 | GOF | GOF | GOF | STK11_p.G242W | Olschwang et al., 2001 | LOF | LOF | LOF |
| KRAS_p.G12C | Prior et al., 2012 | GOF | GOF | GOF | STK11_p.D194Y | Engel et al., 2014 | LOF | LOF | LOF |

**Supplementary Table 2.** Results of the false-positive analysis with mock alleles.

| Control Allele | FPR |
| --- | --- |
| EGFR_WT.c | 7.70% |
| EGFR_p.L858R | 7.80% |
| EGFR_p.T790M, p.L858R.o | 8.00% |
| KEAP1_WT.c | 6.60% |
| KRAS_p.G12V | 7.10% |
| MDM2_WT.c | 5.40% |
| NFE2L2_WT.c | 7.60% |
| STK11_WT.c | 7.90% |
| EMPTY | 2.70% |
| Average | 6.75% |

